# Photosynthesis from stolen chloroplasts increases sea slug reproductive fitness

**DOI:** 10.1101/2021.06.11.448027

**Authors:** Paulo Cartaxana, Felisa Rey, Charlotte LeKieffre, Diana Lopes, Cédric Hubas, Jorge E. Spangenberg, Stéphane Escrig, Bruno Jesus, Gonçalo Calado, Rosário Domingues, Michael Kühl, Ricardo Calado, Anders Meibom, Sónia Cruz

## Abstract

Some sea slugs are able to steal functional chloroplasts (kleptoplasts) from their algal food sources, but the role and relevance of photosynthesis to the animal host remain controversial. While some researchers claim that kleptoplasts are slowly digestible ‘snacks’, others advocate that they enhance the overall fitness of sea slugs much more profoundly. Our analysis show light-dependent incorporation of ^13^C and ^15^N in the albumen gland and gonadal follicles of the sea slug *Elysia timida*, representing translocation of photosynthates to kleptoplast-free reproductive organs. Long-chain polyunsaturated fatty acids with reported roles in reproduction were produced in the sea slug cells using labelled precursors translocated from the kleptoplasts. Finally, we report reduced fecundity of *E. timida* by suppressing kleptoplast photosynthesis. The present study provides the first thorough experimental evidence that photosynthesis enhances the reproductive fitness of kleptoplast-bearing sea slugs, confirming the biological relevance of this remarkable association between a metazoan and an algal-derived organelle.

**Teaser:** Sea slugs incorporate functional chloroplasts from algae and use products of photosynthesis to maximize reproductive output.

## Introduction

Sacoglossa is a group of sap-sucking sea slugs that feed on macroalgae. The most striking feature of some of these sea slugs is their ability to digest the algal cellular content while retaining intact functional chloroplasts (kleptoplasts) within the cells of their digestive gland (1, 2). This process of stealing plastids from algal cells (kleptoplasty) is more common in single-celled eukaryotes, such as foraminiferans, dinoflagellates, and ciliates (3). Recently, Van Steenkiste et al. (4) identified short-term functional kleptoplasts in two species of marine flatworms. However, among metazoans, the capacity for long-term maintenance (up to several months) of functional chloroplasts remains a unique feature of a few species of sacoglossans (5–7). Functional kleptoplasty occurs despite the absence of genetic material with an important role in chloroplast regulation, as these genes have been transferred to the algal nucleus over the evolution of endosymbiosis (8).

The importance of kleptoplasty for the nutrition and metabolism of sacoglossan sea slugs remains controversial. Most studies have shown that photosynthesis plays an important role in individual survival and fitness over periods of food scarcity (9–12), while others argue that it is not essential for slugs to endure starvation (13). Transcriptomic data on the sea slug *Elysia chlorotica* show that chloroplast sequestration leads to significant changes in host gene expression patterns throughout uptake and maturation, similar to that occurring during the establishment of symbiosis in corals, and suggest parallels between these animal–algal interactions (14).

Earlier radiolabeled carbon-based studies indicate translocation of photosynthesis-derived metabolites from functional kleptoplasts into sacoglossan sea slug tissues (15–17). Trench et al. (15) reported ^14^C-labelling within 2 h of incubation in the renopericardium, the cephalic neural tissue and the mucus secreting pedal gland of *Elysia crispata* and *Elysia diomedea*. Recently, Cruz et al. (18) have shown initial light-dependent incorporation of ^13^C and ^15^N in digestive tubules followed by a rapid translocation and accumulation in kleptoplast-free organs of *Elysia viridis*, i.e., in tissues involved in reproductive functions such as the albumen gland and gonadal follicles. However, no direct relation between photosynthesis and reproductive investment of kleptoplast-bearing sea slugs has been established.

In the present study, we investigated the putative role of kleptoplast photosynthesis in the reproduction of the sacoglossan sea slug *Elysia timida* by (i) tracking short-term light-dependent incorporation of inorganic carbon and nitrogen into animal tissues using compound specific isotope analysis (CSIA) of fatty acid methyl esters (FAME) and high-resolution secondary ion mass spectrometry (NanoSIMS), and (ii) investigating the effects of inhibiting photosynthesis (rearing animals under non-actinic light levels) in the number and fatty acid (FA) composition of spawned eggs. We report strong experimental evidence for a role of photosynthesis in the reproductive investment and fitness of a kleptoplast-bearing sea slug.

## Results

### Light-dependent incorporation of C and N

#### NanoSIMS isotopic imaging

Semi-thin section imaging combined with NanoSIMS imaging showed that ^13^C- and ^15^N-labeling was not homogenously distributed in different sea slug tissues, as ^13^C- and ^15^N-hotspots could be observed (Figs. 1–3). NanoSIMS images from individuals incubated in light for 6 to 36 h with ^13^C-bicarbonate and ^15^N-ammonium showed marked ^13^C- and ^15^N-labeling in kleptoplast-bearing digestive tubules (Fig. 1; Supplementary Fig. S1). Individuals incubated in the dark for 36 h displayed no ^13^C-enrichment (Supplementary Fig. S1). In contrast, ^15^N-labeling was observed in the digestive tubules of sea slugs incubated in the dark for 36 h, although at a much lower level than in conspecifics incubated under light (Supplementary Fig. S1).

**Fig. 1.**
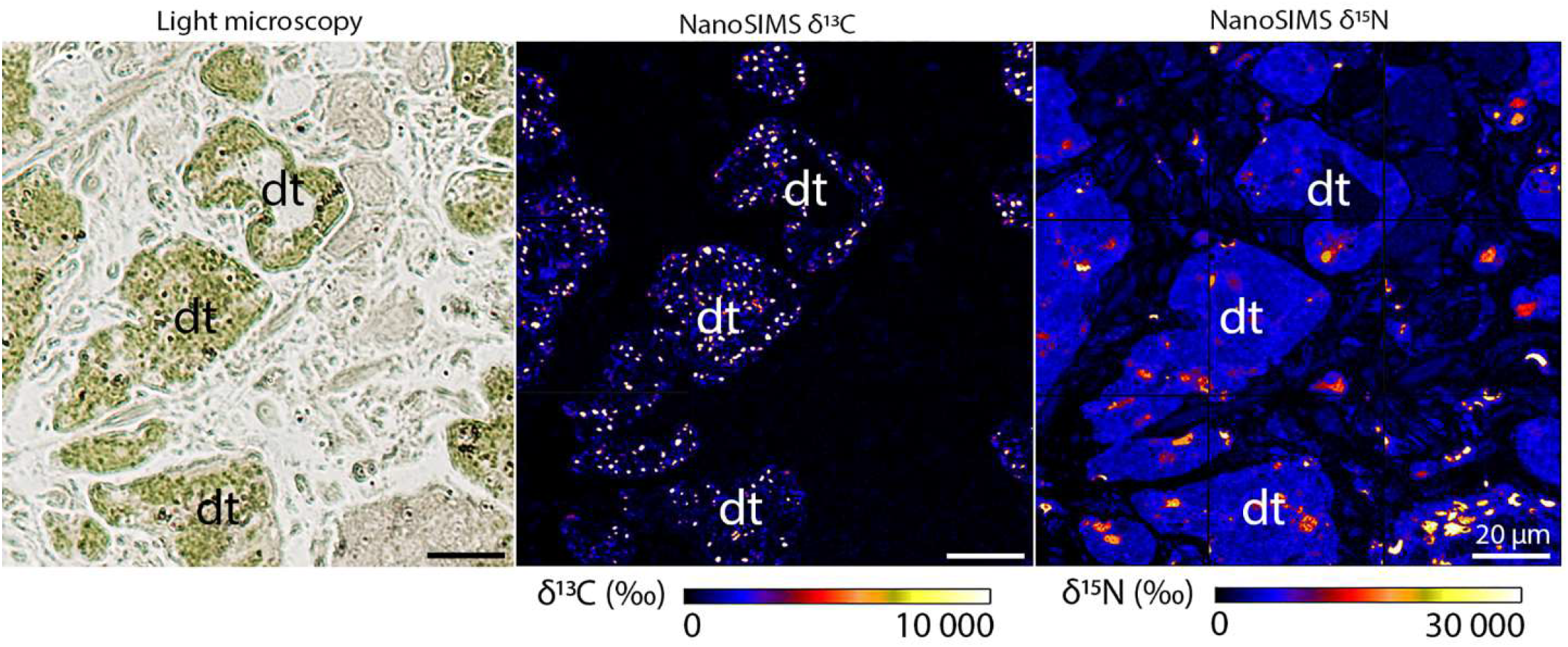
^13^C and ^15^N incorporation in the digestive tubules of *Elysia timida*. Light microscopy picture and corresponding δ^13^C and δ^15^N NanoSIMS images of *E. timida* incubated in artificial seawater enriched with 2 mM NaH^13^CO_3_ and 20 µM ^15^NH_4_Cl, for 6 h in the presence of light; dt – digestive tubules.

Marked ^13^C- and ^15^N-labeling was also observed in the albumen gland and the gonadal follicles (both kleptoplast free) of *E. timida* incubated in the light for 6 to 36 h with ^13^C-bicarbonate and ^15^N-ammonium (Figs. 2 and 3; Supplementary Fig. S1). ^13^C and ^15^N-labeling was still observed in the chasing phase, after individuals were transferred to fresh non-labelled artificial sea water (ASW) for up to another 12 h (Figs. 2 and 3; Supplementary Fig. S1). Again, no ^13^C-enrichment and lower ^15^N-labeling was observed in the albumen gland and the gonadal follicles of sea slugs incubated in the dark for 36 h, when compared to animals incubated in the presence of light (Figs. 2 and 3; Supplementary Fig. S1).

**Fig. 2.**
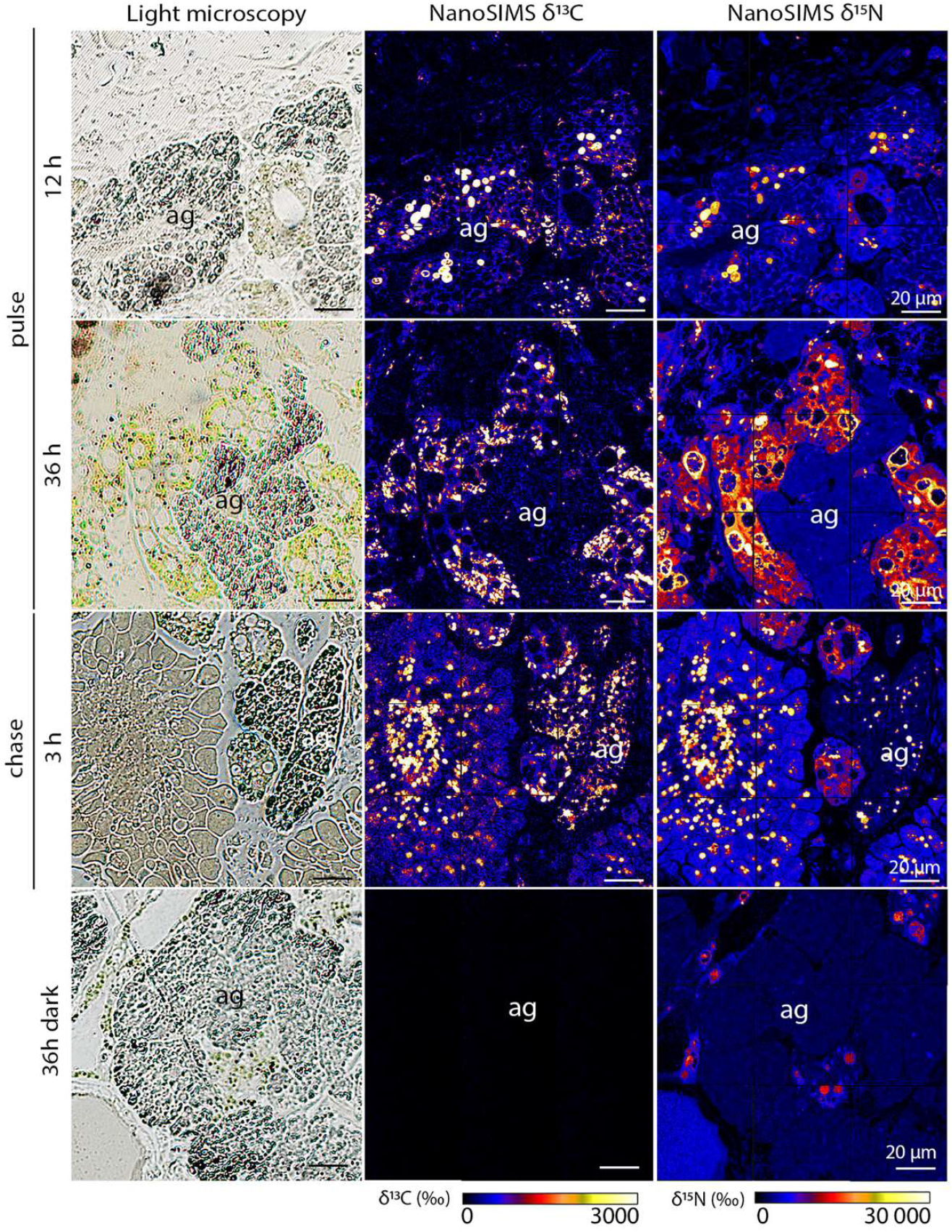
^13^C and ^15^N incorporation in the albumen glands of *Elysia timida*. Light microscopy pictures and corresponding δ^13^C and δ^15^N NanoSIMS images of *E. timida* in an isotopic dual labelling pulse-chase experiment incubated in artificial seawater enriched with 2 mM NaH^13^CO_3_ and 20 µM ^15^NH_4_Cl, in the presence of light for pulse (12 and 36 h) and chase (3 h), and in the dark for 36 h; ag – albumen glands.

**Fig. 3.**
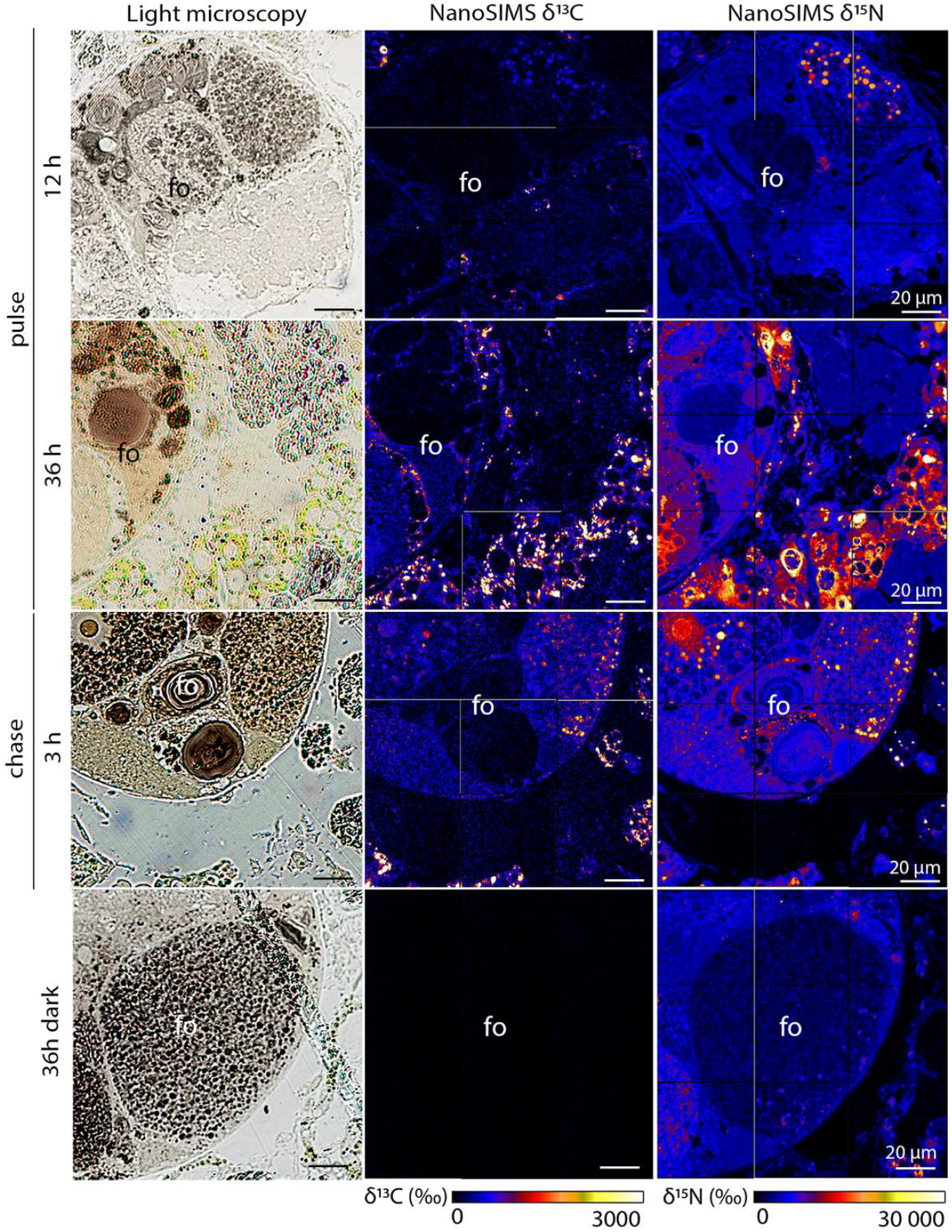
^13^C and ^15^N incorporation in the gonadal follicles of *Elysia timida*. Light microscopy pictures and corresponding δ^13^C and δ^15^N NanoSIMS images of *E. timida* in an isotopic dual labelling pulse-chase experiment incubated in artificial seawater enriched with 2 mM NaH^13^CO_3_ and 20 µM ^15^NH_4_Cl, in the presence of light for pulse (12 and 36 h) and chase (3 h), and in the dark for 36 h.; fo – gonadal follicles.

#### Fatty acid analysis

The most abundant FA (> 5% relative abundance) observed in *E. timida* were the saturated FA (SFA) 16:0 and 18:0, the monounsaturated FA (MUFA) 18:1*n*-9, and the polyunsaturated FA (PUFA) 18:2*n*-6, 18:4*n*-3, 20:4*n*-6, 20:5*n*-3, 22:4*n*-6, and 22:5*n*-3 (Supplementary Table S1). In the presence of light, individuals incubated in ^13^C-bicarbonate enriched ASW for up to 36 h showed an increasing incorporation of ^13^C into FA over time (Fig. 4; Supplementary Table S2). Incorporation of ^13^C occurred in all of the most abundant *E. timida* FA, except 18:4*n*-3 (Fig. 4). Levels of ^13^C labelling decreased from SFA and MUFA precursors to longer-chain PUFAS. However, levels of incorporation of ^13^C in longer-chain FA 22:4*n*-6 and 22:5*n*-3 were higher than in 20:4*n*-6 and 20:5*n*-3, respectively (Fig. 4). In the chasing phase, when individuals were transferred to fresh non-labelled ASW, incorporation of ^13^C into FA generally levelled out (Fig. 4). When animals were kept under dark conditions during 36 h of incubation with ^13^C-bicarbonate enriched ASW, sea slugs’ FA showed no ^13^C-enrichment, with an incorporation equivalent to that of conspecifics incubated in the presence of light but in non-labelled ASW (Fig. 4; Supplementary Table S2).

**Fig. 4.**
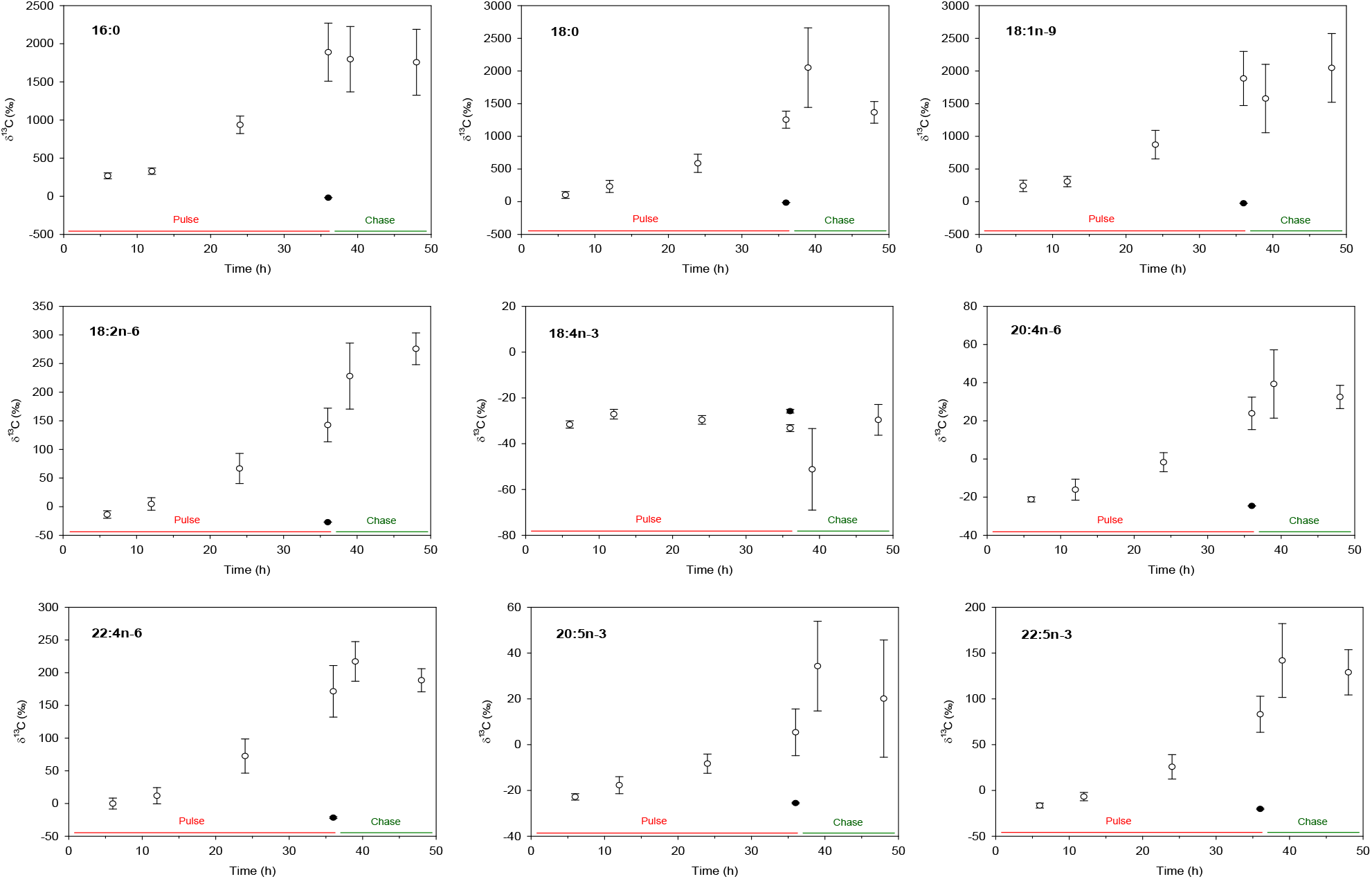
^13^C incorporation in the main fatty acids of *Elysia timida*. ^13^C (‰) in most abundant fatty acids of *E. timida* as a function of time (h) in a isotopic labelling pulse-chase experiment in artificial seawater enriched with 2 mM NaH^13^CO_3_ in the presence of light (open circles, ○) or in dark-incubated specimens for 36 h (closed circles, ●). Mean ± SE, n = 3.

#### Acetabularia acetabulum

FA composition showed a lower diversity to that of *E. timida* (Supplementary Table S1). Lower relative abundances of long-chain PUFA were found in the macroalgal tissue when compared to *E. timida*. PUFA 20:4*n*-6 was present in a relative abundance of 0.26% in the algae compared to 6.15% in the sea slug, while 22:4*n*-6, a major FA in *E. timida* (10.13%), was not present in *A. acetabulum*.

### Effects of light treatment on egg masses

Pairs of *Elysia timida* initiated mating by meeting head-to-head and starting penis protrusion (Supplementary Fig. S2A). Animals mutually inserted their penis into the partner’s female aperture, located at the base of the right parapodium. Spiral-shaped egg masses (Supplementary Fig. S2B) were spawned by *E. timida* in both light and quasi-dark conditions, although light treatment affected the number of spawning events. Sea slugs reared in light (40-160 µmol photons m^−2^ s^−1^) produced 7.5±0.2 egg masses per pair (mean±std error), corresponding to 7 or 8 egg masses over the 28-days experimental period. Spawning activity was more variable in pairs reared under low non-actinic light levels (5 µmol photons m^−2^ s^−1^) and ranged from 1 to 5 egg masses per pair (3.0±0.7, mean±std error). The number of eggs produced by animals reared in light was significantly higher (*t*_9_ = 3.521, p = 0.007) than for sea slugs reared under quasi-dark conditions (238±13 vs 129±30 eggs slug^−1^ week^−1^, respectively; Fig. 5). FA concentrations per egg were not significantly affected by light treatments (Fig. 6; Supplementary Table S3). FA composition was similar in *E. timida* individuals and egg masses (Supplementary Tables S1 and S3).

**Fig. 5.**
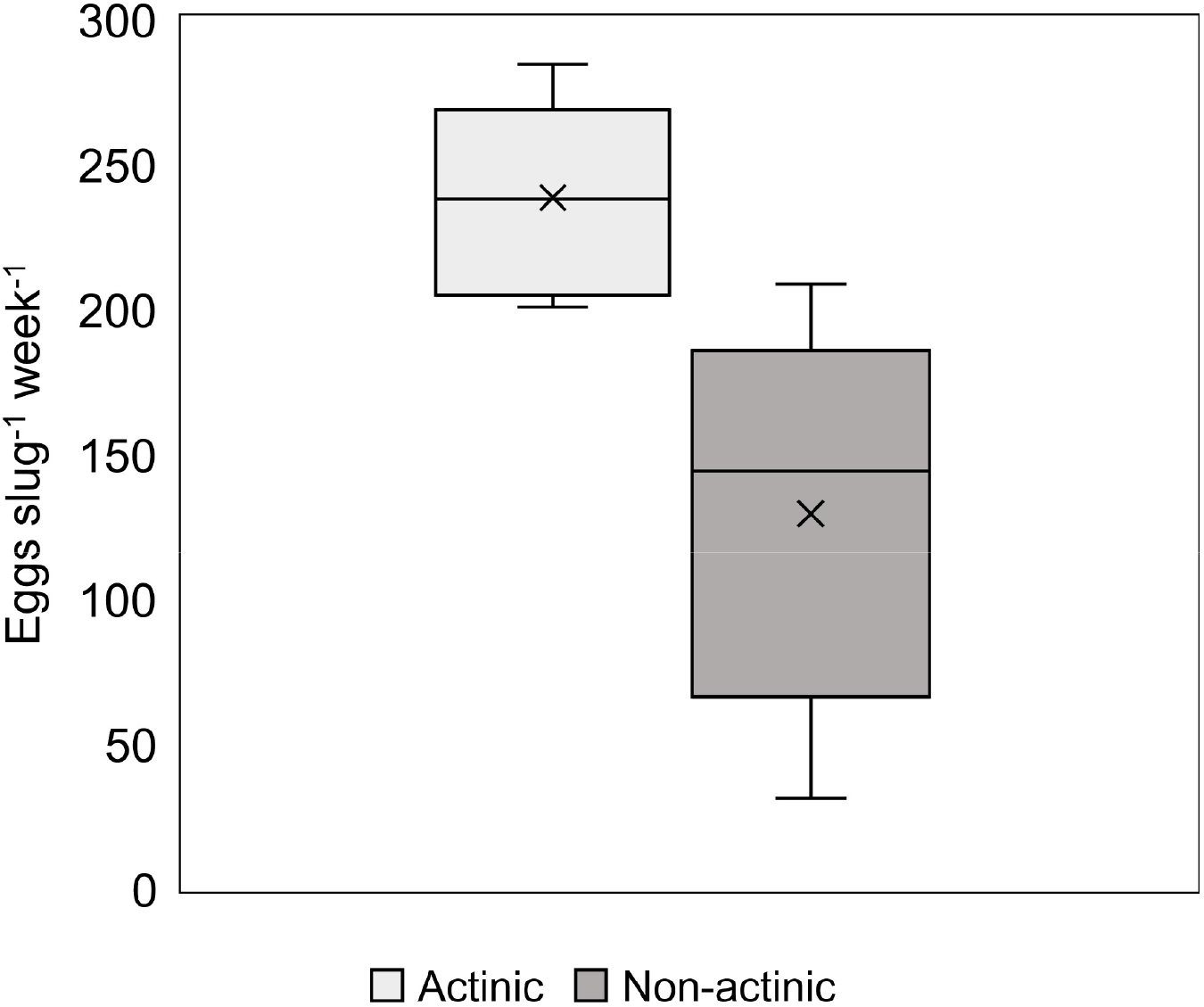
Fecundity of *Elysia timida*. Number of eggs spawned by *E. timida* exposed to a 14:10 h light/dark photoperiod and a scalar irradiance of 40-160 µmol photons m^−2^ s^−1^ (Actinic) or 5 µmol photons m^−2^ s^−1^, (Non-actinic) for 28 days. The line is the median, the x represents the mean, top and bottom of the box are the 75% and 25% percentile, and the whiskers represent the maximum and minimum values. Animals were fed continuously with *Acetabularia acetabulum*. Differences between treatments were significant at p < 0.007.

**Fig. 6.**
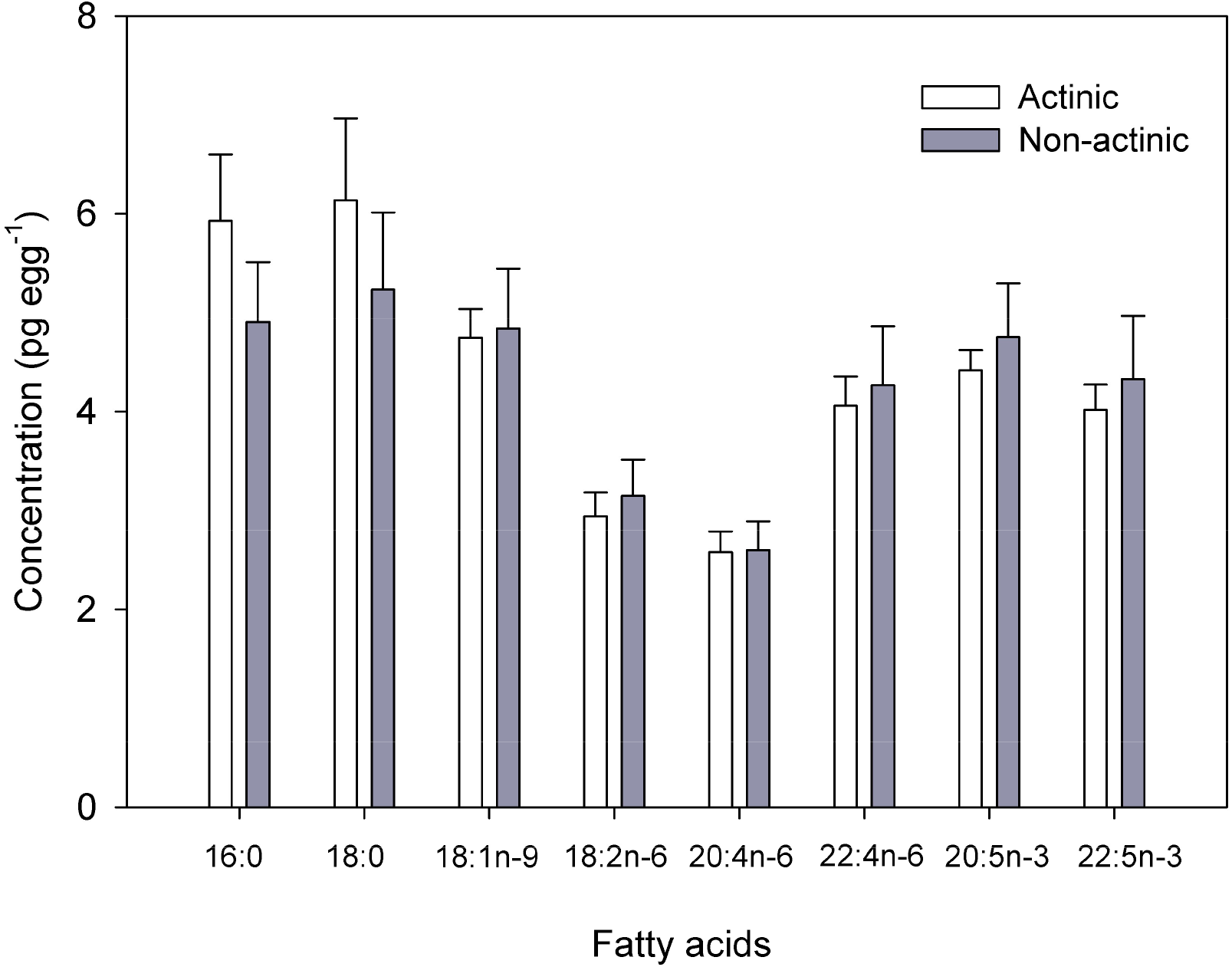
Most abundant fatty acids in the egg masses of *Elysia timida*. Concentration of most abundant fatty acids in egg masses spawned by *E. timida* (mean ± SE, n = 6 or 5) exposed to a 14:10 h light/dark photoperiod and a scalar irradiance of 40-160 µmol photons m^−2^ s^−1^ (Actinic) or 5 µmol photons m^−2^ s^−1^ (Non-actinic) for 28 days. The egg masses analyzed were the last one spawned by sea slug pairs stocked on each experimental unit. Animals were continuously fed with *Acetabularia acetabulum*.

## Discussion

NanoSIMS isotopic imaging of the sea slug *E. timida* showed inorganic ^13^C incorporation in kleptoplast-bearing digestive tubules of light exposed animals. However, light-dependent carbon incorporation was not restricted to these kleptoplast-bearing cells, and rapid accumulation (within 6 h) was observed in kleptoplast-free organs such as the albumen gland and gonadal follicles. ^13^C incorporation was not detected in the tissues of *E. timida* exposed to full darkness. Thus, our data clearly demonstrate that inorganic ^13^C was photosynthetically accumulated into functional kleptoplasts and subsequently translocated to other sea slug tissues, likely through soluble C-compounds (e.g. sugars) or FA.

Translocation of photosynthetically acquired carbon to animal tissues was previously identified in other species of sacoglossan sea slugs: *Elysia crispata, Elysia diomedea, Elysia viridis* and *Plakobranchus ocellatus* (15–18). Using ^14^C, Trench et al. (15) observed labelling of kleptoplast-bearing digestive tubules after 15 min of light incubation for *E. crispata* and *E. diomedea*. Carbon incorporation was also detected in kleptoplast-free organs, such as the renopericardium (after 1 h), the cephalic neural tissue and the mucus secreting pedal gland (after 2 h), and the intestine (after 5 h) (15). Trench et al. (16) observed that ^14^C was incorporated into glucose and galactose in *E. viridis*, while Ireland & Scheuer (17) reported carbon incorporation in sugars and polypropionates for *P. ocellatus*. Using electron microscopy combined with NanoSIMS imaging, Cruz et al. (18) observed ^13^C-labeling after 1.5 h in starch grains of kleptoplasts present in the kleptoplast-bearing digestive tubules of *E. viridis*, but ^13^C-labeling was also found in the cytoplasm surrounding the photosynthetic organelles. After longer incubation times (1.5 to 12 h), ^13^C-labeling was detected in *E. viridis* organs involved in reproduction, namely the albumen gland and gonadal follicles (18). Evidence of fast translocation of photosynthates to kleptoplast-free animal tissues is not compatible with a previously proposed hypothesis that kleptoplasts are slowly digestible food reserves and that photosynthates produced are not continuously made available to the slug (13, 19).

Sea slugs showed a much higher level of ^15^N-enrichment in their tissues when incubated in the presence of light. Light-dependent incorporation of ^15^N was previously reported for *E. viridis* (18, 20). Teugels et al. (20) identified glutamine synthetase (GS) – glutamate synthetase (GOGAT) as the main pathway involved in N incorporation in the kleptoplasts. Hence, kleptoplasts may not only provide energy and carbon skeletons, but could also play a role in protein synthesis. *De novo* protein synthesis has been shown to occur for plastid-encoded membrane proteins in *Elysia chlorotica*, even after several months of starvation (21). Contrary to ^13^C, our NanoSIMS imaging of *E. timida* recorded ^15^N-incorporation in the dark. Nitrogen incorporated in specimens incubated in the dark (albeit significantly reduced) could result from the glutamine dehydrogenase (GDH) pathway in mitochondria (18, 20).

The labelling with ^13^C was done in the absence of *A. acetabulum*, the macroalgal food source of *E. timida*, safeguarding that the labelled FA detected were not obtained heterotrophically (i.e., trough grazing on *A. acetabulum*). Instead, ^13^C-labelled FA must have been synthesized in the kleptoplasts of the digestive tubules and eventually translocated to other animal cells. Photosynthetic lipid production has been suggested to play an important role in the establishment of kleptoplasty in photosynthetic sea slugs (22, 23). Additionally, labelled FA could have been produced in the animal cells through elongation/desaturation reactions using labelled precursors translocated from the kleptoplasts. In fact, the presence of labeled 22:4*n*-6 in *E. timida* is a direct evidence that the latter process occurred, as this FA was not present in *A. acetabulum*.

It was generally assumed that animals were unable to biosynthesize PUFA *de novo* since, presumably, they lacked specific desaturases required to produce 18:2*n*-6 (LA; linoleic acid) (24). However, several findings have challenged this long-held assumption and it was recently shown that Δ desaturases genes enabling *de novo* PUFA biosynthesis are widespread among invertebrates (25). *De novo* biosynthesis of PUFA can occur via different pathways (26). Tracking of ^13^C-labelled FA allowed us to infer the main pathway of PUFA biosynthesis in *E. timida* (Fig. 7). A Δ9 desaturase likely mediated the insertion of the first unsaturation in 18:0 (stearic acid) to produce oleic acid (OA; 18:1n-9). The introduction of further unsaturations into OA must have proceeded via a pathway involving Δ12 desaturases to produce LA, which was subsequently desaturated to 18:3*n*-3 (ALA; α-linolenic acid) by the action of a Δ15 desaturase. However, we observed limited ^13^C incorporation in ALA (δ^13^C = −6.6 ‰ after 36 h) and no incorporation into 18:4*n*-3 or 18:3*n*-6 FA (δ^13^C = −33.2 and −38.4 ‰ after 36 h, respectively) (Supplementary Table S2). This suggests that Δ6 desaturases enabling the production of 20:3*n*-6 and 20:4*n*-3 from LA and ALA, respectively, are likely absent in *E. timida* (Fig. 7). PUFA biosynthesis probably proceeded via alternative elongase → Δ8 desaturase-mediated reactions from LA, as suggested by the presence of ^13^C-labelled 20:2*n*-6 and 20:3*n*-6 FA (δ^13^C = +191.4 and +52.4 ‰ after 36 h, respectively) (Supplementary Table S2). The fatty acid 20:5*n*-3 (eicosapentaenoic acid, EPA) was likely produced from 20:3*n*-6 by the action of Δ17 → Δ5 desaturases and 22:5*n*-3 from EPA by an elongase mediated reaction. Long-chain PUFA 20:4*n*-6 (arachidonic acid; ARA) was most probably synthesized from 20:3*n*-6 by the action of a Δ5 desaturase, with a further elongase enabling 22:4*n*-6 production (Fig. 7).

**Fig. 7.**
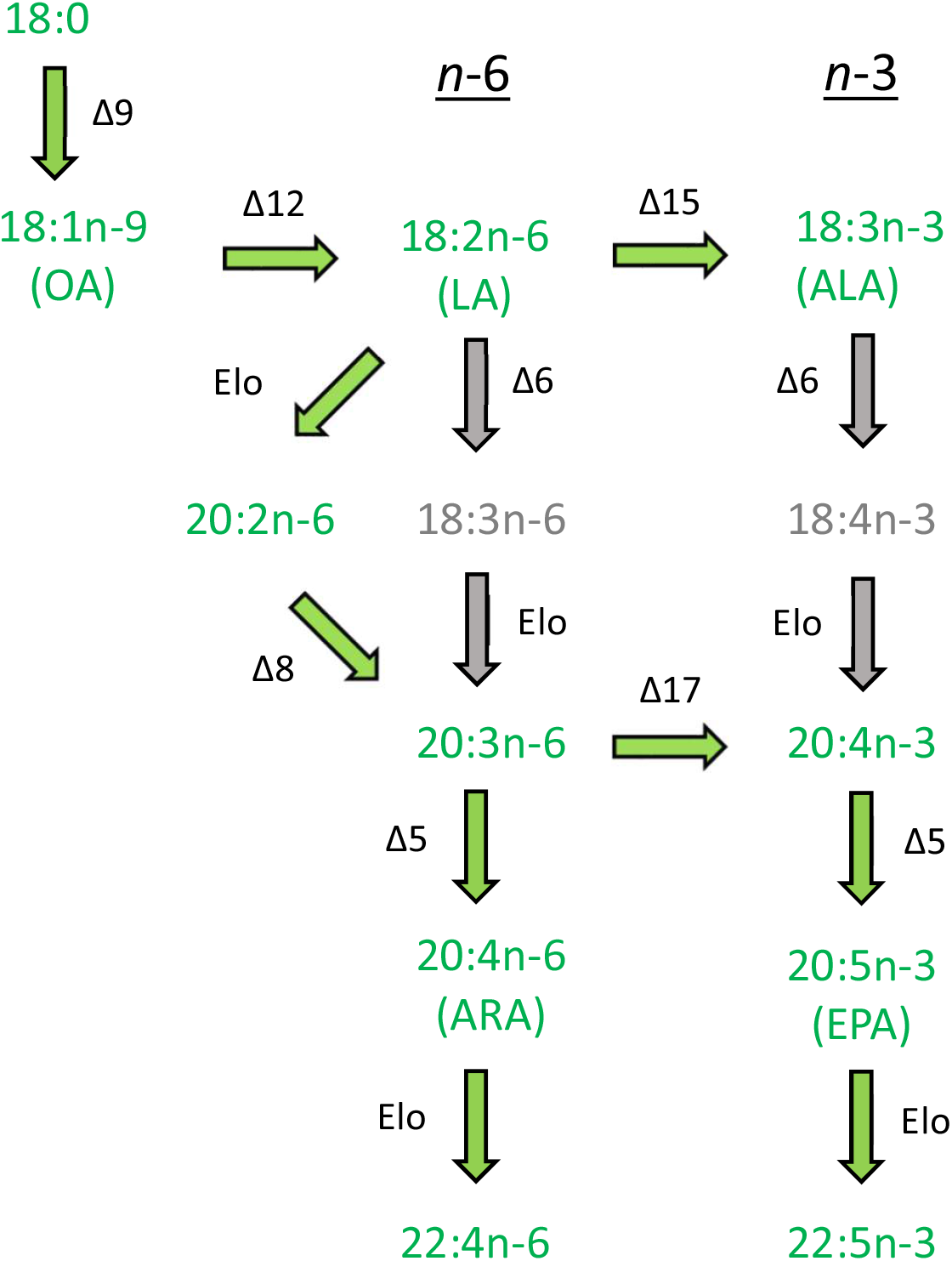
Biosynthetic pathway of polyunsaturated fatty acids in *Elysia timida*. Fatty acids in green showed light-dependent ^13^C incorporation. Fatty acids in grey showed no ^13^C incorporation. Desaturase enzymes are denoted with “Δ” and elongases with “Elo”. OA: oleic acid, LA: linoleic acid, ALA: α-linolenic acid, ARA: arachidonic acid, EPA: eicosapentaenoic acid.

A general dilution of the ^13^C signal was observed along the FA biosynthetic pathway from saturated and monounsaturated precursors 18:0 and OA to longer-chain PUFA. However, an increase in the ^13^C signal was observed in the last elongation steps of 22:5*n*-3 and 22:4*n*-6 production from EPA and ARA, respectively. This finding indicates that the carbon donor (malonyl-CoA) during this elongation process was ^13^C-enriched and thereby preferentially provided by kleptoplasts. Torres et al. (27) reported that methylmalonyl-CoA incorporating kleptoplast fixed-carbon is used by sacoglossan sea slugs in the synthesis of UV- and oxidation-blocking polypropionate pyrones by the action of FA synthase-like proteins. Pyrones could be critical for maintaining long-term photosynthetic activity in sacoglossan sea slugs by serving antioxidant and photoprotective roles (17, 27).

Dietary PUFA have been shown to modulate marine invertebrate gametogenesis, embryogenesis and larval development (28–30). The levels of PUFA recorded in female tissues and embryos of the sea snail *Crepidula fornicata* were related to its reproductive output (31). Bautista-Teruel et al. (32) reported that the reproductive performance in the gastropod *Haliotis asinina* was linked to diets with increased levels of PUFA, such as EPA and ARA. The latter fatty acids are precursors of prostaglandins, a group of biologically active compounds participating in marine invertebrate reproduction (33, 34). Hence, the assembly of photosynthesis-driven long-chain PUFA as shown in our study for *E. timida* (in the kleptoplasts and elsewhere in animal cells using photosynthesis-derived precursors) are likely to play a crucial role in the reproductive output of this species and increase evolutionary fitness.

Reproductive investment, assessed as the number of eggs spawned by *E. timida* along a 4 week-period, was significantly higher in actinic light-exposed sea slugs (40-160 µmol photons m^−2^ s^−1^) than in animals reared under non-actinic conditions (5 µmol photons m^−2^ s^−1^). Under a period of resource shortage (i.e. inhibited photosynthesis under non-actinic light levels), *E. timida* clearly reduced its reproductive energy investment by decreasing the number of spawned egg masses. Shiroyama et al. (35) observed higher number of eggs spawned by *Elysia atroviridis* when fed under light (30 µmol photons m^−2^ s^−1^) than when kept under non-actinic conditions (1 µmol photons m^−2^ s^−1^). The availability of the macroalga *Codium tomentosum* was also shown to affect the number of eggs spawned by the sea slug *E. viridis* (36). However, in the latter case, limited access to food affected mainly the number of eggs per egg mass rather than the number of spawning events.

In our study, complete darkness was not used to inhibit photosynthesis in order not to disrupt diel biorhythms and animal behaviour. The sea slug *E. viridis* was reported to become inactive under full darkness, rarely seeming to feed (9). Hence, an extremely low light intensity (i.e. non-actinic light), enough to substantially inhibit kleptoplast photosynthesis, was used instead of full darkness (37). Animals in both light treatments were observed attached to the macroalgae, and the cellular content of *A. acetabulum* was emptied similarly in actinic and non-actinic light treatments. This ensured that the heterotrophic feeding ability of sea slugs was not affected by the experimental conditions, and it is thus reasonable to attribute the differences in fecundity to the resources provided by kleptoplast photosynthesis. Although reproductive investment was reduced when photosynthesis was inhibited, *E. timida* allocated similar amounts of FA to individual eggs regardless of light treatment. This is particularly relevant for egg viability in species such as *E. timida*, in which offspring success depends exclusively on the parental provisioning of endogenous reserves to fuel embryonic development and early larval life, until lecithotrophic larvae are able to metamorphose to imago of the adult and feed on exogenous food sources (38).

In conclusion, we report the allocation of photosynthates to kleptoplast-free organs involved in the reproduction of *E. timida*, along with photosynthesis-driven assembly of long-chain PUFA, and higher sea slug fecundity under actinic light conditions. These results demonstrate that kleptoplast photosynthesis increases reproductive investment of *E. timida*. It has been shown that kleptoplasty in Sacoglossa contributes to survival and fitness in periods of food scarcity, in some cases allowing individuals to endure several months of starvation (9, 10, 39). We show that functional kleptoplasty in sacoglossan sea slugs may further potentiate species’ success by maximizing its reproductive output.

## Materials and Methods

### Animal collection and maintenance

Specimens of *Elysia timida* (Risso, 1818) (Supplementary Fig. S2A) were collected in Puerto de Mazarrón in the Mediterranean Sea, Spain. Sampling of *E. timida* and its algal food, *Acetabularia acetabulum* (Linnaeus) P. C. Silva, 1952, was done by SCUBA diving at a depth of *∼*2 m. Animals and macroalgae were kept in aerated seawater collected at the sampling site and transported to the laboratory within 48 h. Sea slugs and macroalgae were maintained for 2 weeks in a 150-L recirculated life support system (LSS) operated with ASW at 18°C and a salinity of 35. The photoperiod was kept at 14 h light:10 h dark, with a photon scalar irradiance of 60 µmol photons m^−2^ s^−1^ being provided by T5 fluorescent lamps. Photon scalar irradiance was measured with a Spherical Micro Quantum Sensor and a ULM–500 Universal Light Meter (Heinz Walz GmbH, Germany). The laboratory adaptation period was chosen to ensure replicability in feeding and light history of the animals at the beginning of the experiment.

### Light-dependent incorporation of C and N

#### Dual isotopic labelling incubations

Isotopic dual labelling pulse-chase experiments were conducted in closed-systems (1-L glass bottles, 3 independent containers per treatment). Labelled ASW was prepared in accordance with Harrison et al. (40), but using NaH^13^CO_3_ (^13^C isotopic abundance of 99%, Sigma-Aldrich) and ^15^NH_4_Cl (^15^N isotopic abundance of 98%, Sigma-Aldrich) to a final concentration of 2 mM and 20 µM, respectively (labelled-ASW). Non-labelled ASW (control-ASW) contained NaHCO_3_ and NH_4_Cl (Sigma-Aldrich) in the same concentrations as the isotopically enriched-ASW. Sea slugs were incubated in labelled- and control-ASW at 18ºC, in the absence of their food source, and under a photon scalar irradiance of 100 µmol photons m^−2^ s^−1^ (measured inside the glass bottles). Additionally, sea slugs in labelled-ASW were incubated in full darkness (Supplementary Table S4). Dark conditions served as a control for light-independent carbon and nitrogen incorporation. The pulse of isotopic dual labelling started 1 h after the onset of the light period. A subset of three individuals kept in labelled-ASW and exposed to light were sampled after 6, 12, 24 and 36 h of incubation (pulse phase), quickly rinsed with distilled water, flash frozen in liquid nitrogen and stored at –80ºC until further FA analysis. An additional individual kept in labelled-ASW and exposed to light was collected at each of the referred time points, rinsed and fixed in 0.2 M cacodylate buffer containing 4% glutaraldehyde and 0.5 M sucrose and stored at 4ºC for 24 h before tissue preparation for secondary ion mass spectrometry imaging (NanoSIMS 50L). Individuals from labelled-ASW incubated in dark conditions and from control-ASW exposed to light were sampled after 36 h as described above for FA and SIMS analysis. Remaining individuals in labelled-ASW and exposed to light were transferred to fresh control-ASW. During this chase period, a subset of individuals was collected after 3 and 12 h as described above, for FA and SIMS analysis (Supplementary Table S4). In light treatments, light was set constant throughout the 48 h experiment.

#### Tissue preparation for NanoSIMS imaging

Sea slugs kept in the fixative for 24 h at 4ºC were transferred to 0.2 M cacodylate buffer with decreasing sucrose concentrations (15 min in cacodylate buffer with 0.5 M sucrose, then 0.25 M sucrose and finally no sucrose) and finally transferred to 2% osmium tetroxide in distilled water for 1 h at room temperature in the dark. Sea slugs were then dehydrated in an increasing series of ethanol concentrations (two times 10 min in 30, 50, 70, 90 and 96% ethanol and two times 20 min in 100% ethanol; room temperature) followed by two times 10 min in acetone before resin embedding. Sea slugs were transferred to acetone:epon resin (1:1) overnight before being fully embedded in 100% epon resin for 6 h in a turning wheel. Sea slugs were finally transferred to new 100% epon resin and dried at 60°C for 48 h. Overview semi-thin cuts of 1.5 µm thickness were made from the sea slug body part roughly after the pericardium. Semi-thin sections were cut on a Leica UC7 ultramicrotome using a Leica glass knife and were placed on circular glass cover slips. Histological overviews were documented on an optical light microscope. Before NanoSIMS analysis, semi-thin sections were coated with a ca. 15 nm thick gold layer.

#### High-resolution secondary ion mass spectrometry (NanoSIMS) isotopic imaging

Large areas of interest were imaged with a NanoSIMS 50L secondary ion mass spectrometer. This allowed imaging of the subcellular distribution of ^13^C and ^15^N enrichment in the exact same areas of the imaged histological overviews described above, enabling a direct correlation of structural and isotopic images. All measurements were performed using the following analytical conditions: 16 keV primary ion beam of Cs^+^ focused to a beam spot of ca. 100–150 nm and counting ^12^C_2-_, ^13^C^12^C^−, 14^N^12^C^−^ and ^15^N^12^C^−^ ions in electron multipliers at a mass resolution of > 8000 (Cameca definition), enough to resolve potential interferences in the mass spectra. Images captured with NanoSIMS 50L were processed using the L’IMAGE® software (Larry R Nittler, Carnegie Institution of Washington, Washington DC, USA). Regions of interest selecting individual anatomic structures were defined, and distribution maps of ^13^C/^12^C and ^15^N/^14^N ratios were obtained by taking the ratio between the drift-corrected ^13^C^12^C^−^ and ^12^C^12^C^−^ images, and ^15^N^12^C^−^ and ^14^N^12^C^−^ images, respectively. Five stacked planes were used for each image. ^13^C and ^15^N enrichment values in the figures were expressed as delta notations, δ^13^C = (C_mes_/ C_nat_ – 1)*1000 and δ^15^N = (N_mes_/N_nat_ – 1)*1000, where *C*_*mes*_ and *N*_*mes*_ are the measured ^12^C^13^C^−^/^12^C_2_^−^ and ^15^N^12^C^−^/^14^N^12^C^−^ ratios of the sample and *C*_*nat*_ and *N*_*nat*_ is the average ^12^C^13^C^−^/^12^C_2_^−^ and ^15^N^12^C^−^/^14^N^12^C^−^ ratios measured in control, non-labelled samples. A number of measurements on these controls (n=12) yielded distributions of δ^13^C = 0 ± 9.6 ‰, and δ^15^N = 0 ± 21.6 ‰ (±2σ).

#### Compound Specific Isotope Analysis (CSIA) of Fatty Acid Methyl Esters (FAME)

Fatty acid extraction was performed following the method of Bligh and Dyer (41) as modified by Meziane and Tsuchiya (42) and Passarelli et al. (43). Before extraction, an internal standard (C23:0) was added to every sample for quantification purposes (0.5 mg/mL). Lipids were extracted with a 20 min ultrasonication (sonication bath, 80 kHz, Fisherbrand™) in a mixture of distilled water, chloroform and methanol in ratio 1:1:2 (v:v:v, in mL). Lipids were concentrated under N_2_ flux, and saponified, in order to separate FA, with a mixture of NaOH (2 M) and methanol (1:2, v:v, in mL) at 90 °C during 90 min. Saponification was stopped with 500 µL hydrochloric acid. Samples were then incubated with BF_3_-methanol at 90°C during 10 min to transform free fatty acids into fatty acids methyl esters (FAME), which were isolated and kept frozen in chloroform. Just before analysis, samples were dried under N_2_ flux and transferred to hexane. FAME Peaks were identified by comparison of the retention time with analytical standards (Supelco 37 Component FAME Mix, Sigma-Aldrich, Buchs, Switzerland). Additional identification of the samples was performed using a gas chromatograph coupled to mass spectrometer (GC-MS, Varian 450GC with Varian 220-MS). Compounds annotation was performed by comparing mass spectra with NIST 2017 library.

The compound specific isotope analysis of the FAME was performed by gas chromatograph/combustion/isotope ratio mass spectrometry (GC/C/IRMS) with an Agilent 6890 GC instrument coupled to a Thermo Fisher Scientific (Bremen, Germany) Delta V Plus IRMS instrument via a combustion interface III under a continuous helium flow. The GC separation was performed with the HP-FFAP column (50 m × 0.20 mm; length × inner diameter) coated with 0.33 µm nitroterephthalic acid modified polyethylene glycol stationary phase. The FAME samples were injected splitless at 230 °C. After an initial period of 2 min at 100°C, the column was heated to 240°C (held 26 min) at 5°C/min, then to 245°C (held 4 min). This GC conditions were optimized for good separation of unsaturated FAs by injection of a standard mixture of 37 FAMEs (Supelco 37 Component FAME Mix, Sigma-Aldrich, Buchs, Switzerland) containing C4–C24 homologues. For calibration and normalization of the measured FAME δ^13^C values were used the previously determined δ^13^C values (by elemental analysis/IRMS) of a mixture of deuterated carboxylic acids used as external standards. For quality control, the repeatability and intermediate precision of the GC/C/IRMS analysis and the performance of the GC and combustion interface were evaluated every 5 runs by injection of a carefully prepared mixture of FAMEs reference materials (44). The standard deviation for repeatability of the δ^13^C values ranged between ±0.05 and ±0.5 ‰ for m/z 45 peak size between 15000 mV and <500 mV. The FA δ^13^C were determined from the FAME δ^13^C by correction for the isotopic shift due to the carbon introduced by methylation using a mass balance equation (45).

### Effects of light treatment on egg masses

A floating tray with wells (56 mm diameter x 60 mm depth) was placed floating in the described LSS. The bottom of the wells was made of a 0.5 mm-mesh to allow water exchange (36). A re-circulating water pump was placed below the experimental tray to increase water renewal inside the wells. Twenty-four adult *E. timida* specimens were randomly divided in pairs and placed in individual wells. The photoperiod was kept at 14 h light:10 h dark. Two treatments (6 replicates per treatment, each replicate being a pair of sea slugs) were performed: 1) “Actinic” treatment in which the sea slug specimens were subjected to a photon scalar irradiance of 40-160 µmol photons m^−2^ s^−1^, depending on the position inside the well; 2) “Non-actinic” in which the sea slug specimens were subjected to a photon scalar irradiance of 5 µmol photons m^−2^ s^−1^ (non-actinic light level). Light treatments were achieved by placing either transparent or opaque lids over the wells. In the case of the non-actinic treatment, light reached the animals through the bottom mesh. Animals were fed every day with *A. acetabulum* grown at a photon scalar irradiance of 60 µmol photons m^−2^ s^−1^ under a 14 h light:10 h dark photoperiod. During the experimental period, 1 animal died in the non-actinic conditions, reducing the number of replicates in this treatment to n = 5.

*Elysia timida* is a simultaneous hermaphrodite, each individual possessing both male and female sexual systems and with a high degree of synchrony and reciprocity in sperm transfer (46). Egg masses spawned by the sea slugs on the walls of the wells and, occasionally, on the net at the bottom of the wells were counted daily for 28 days and collected using a scalpel (Supplementary Fig. S2B). The number of eggs in each individual egg mass was counted using a Leica DMS300 digital microscope. Egg masses were gently washed in ultrapure water, frozen at −80 °C and freeze-dried. The last egg mass produced in each experimental unit (well) was analyzed for FA composition.

Total lipid extracts from *E. timida* egg masses were extracted using a solid-liquid extraction. Freeze-dried samples were macerated and homogenized with 400 µL of methanol and 200 µL of dichloromethane, sonicated for 1 min and incubated on ice for 30 min on an orbital shaker. An additional volume of dichloromethane (200 µL) was added, followed by centrifugation at 2000 rpm for 10 min. The liquid phase was collected in a new tube, dried under a nitrogen stream and preserved at −20ºC for FA analysis. Five replicates of *E. timida* sea slugs reared in actinic light conditions and *A. acetabulum* were similarly washed in ultrapure water, frozen at −80°C, freeze-dried and macerated prior to lipid extraction. Total lipid extracts of *E. timida* and *A. acetabulum* samples were obtained using the modified method of Bligh and Dyer (41). Briefly, freeze-dried samples were vigorous homogenised with methanol/dichloromethane (600 µL / 300 µL in *E. timida*; 2.5 mL / 1.25 mL in *A. acetabulum*). Samples were sonicated for 1 min, incubated on ice (30 min in *E. viridis*; 2h 30 min in *A. acetabulum*) on an orbital shaker and centrifuged at 2000 rpm for 10 min. The organic phase was collected in a new tube and mixed with dichloromethane and ultrapure water (300 µL/300 µL in *E. timida*; 1.25 mL/2.25 mL in *A. acetabulum*). After centrifugation at 2000 rpm for 10 min the organic phase was collected in a new tube and the aqueous phase was reextracted with dichloromethane (300 µL in *E. timida*; 2 mL in *A. acetabulum*). Both organic phases were dried under a nitrogen stream and preserved at −20ºC until further analysis.

Fatty acids in lipid extracts from the three biological matrices surveyed (egg masses, sea slugs and macroalgae) were transmethylated according to Aued-Pimentel et al. (47) to obtain FAME and analyzed by gas chromatography – mass spectrometry (GC-MS). FAME identification was performed by comparing retention times and mass spectra with those of commercial FAME standards (Supelco 37 Component FAME Mix, ref. 47885-U, Sigma-Aldrich) and confirmed by comparison with the Wiley library and the spectral library from ‘The Lipid Web’ (48). FA quantification was performed using calibration curves obtained from FAME standards under the same instrumental conditions. FA in *E. timida* and *A. acetabulum* were expressed as relative abundances (%). FA concentrations in the eggs were expressed as pg egg^−1^ dividing the FA content of the whole egg mass by the number of eggs.

### Statistical analyses

The number of eggs spawned in each experimental unit (pairs of sea slugs placed on each well) was averaged to avoid pseudo-replication, and averages were treated as independent replicates (49). Statistically significant differences in the number and FA concentrations of eggs spawned by Actinic versus Non-actinic reared animals were tested using independent samples *t*-tests. Normality was checked using a Shapiro-Wilk’s test, while homogeneity of variances was tested using Levene’s test. Statistical analyses were carried out using IBM SPSS Statistics 24.

## Acknowledgments

We thank Dr. José Templado and Dr. Marta Calvo for help in the collection of *E. timida* and *A. acetabulum* and Sofie Jakobsen and Gabriel Ferreira for technical assistance.

## Author contributions

Conceptualization: PC, CH, BJ, GC, MK, RC, AM, SC

Methodology: JS, RD, AM, SC

Investigation: PC, FR, CL, DL, CH, JS, SE, BJ, SC

Supervision: SC

Writing—original draft: PC, SC

Writing—review & editing: All authors

