## Supplementary Figs. S1-S2 & Tables S1-S4 for "Photosynthesis from stolen chloroplasts increases sea slug reproductive fitness"

**This PDF file includes:**

Figs. S1 to S2
Tables S1 to S4

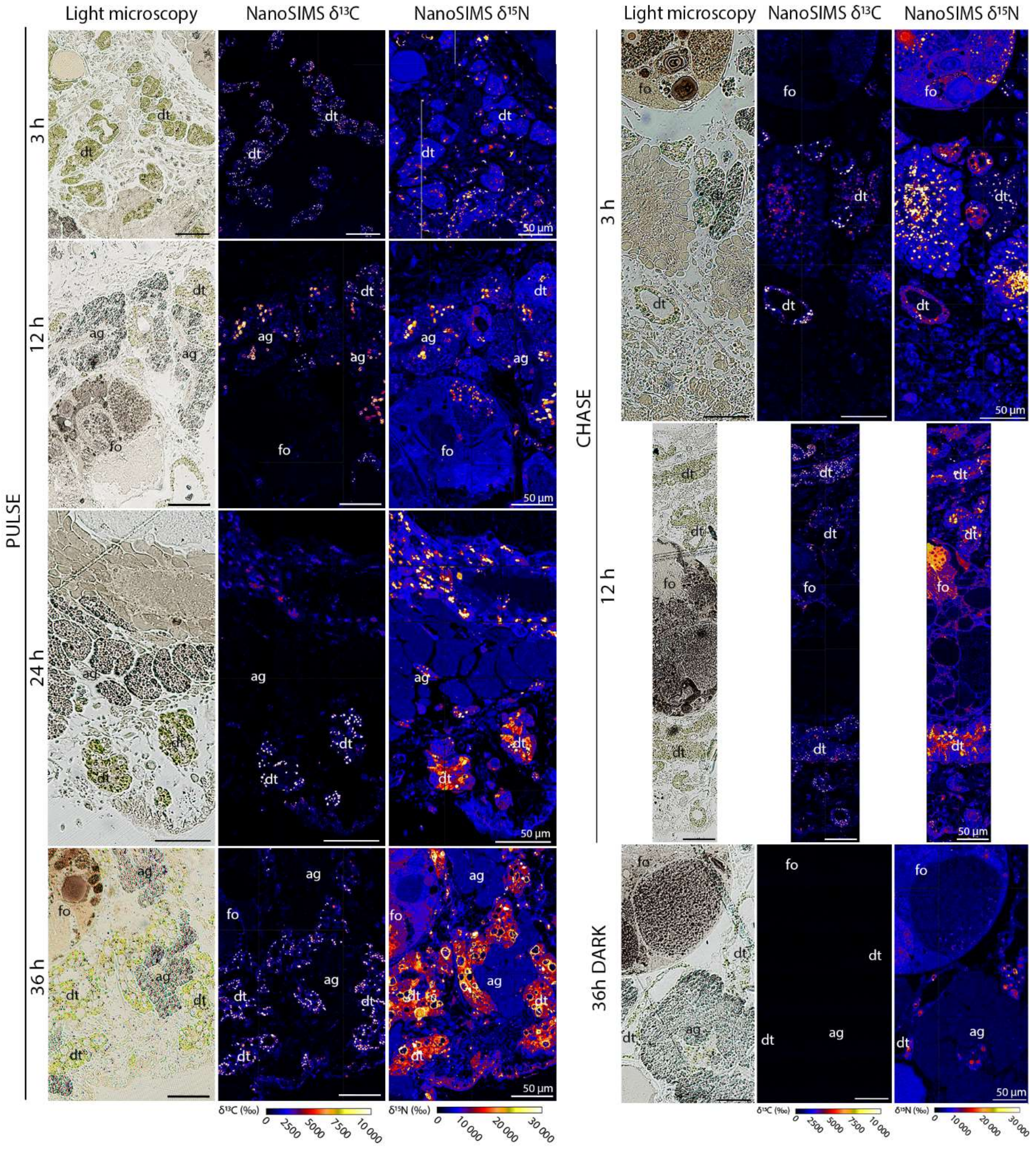


**Fig. S1. $^{13}C$ and $^{15}N$ incorporation in the different tissues of *Elysia timida*.** Light microscopy pictures and corresponding $\delta^{13}C$ and $\delta^{15}N$ NanoSIMS images of *E. timida* in an isotopic dual labelling pulse-chase experiment incubated in artificial seawater enriched with 2 mM $NaH^{13}CO_3$ and 20 µM $^{15}NH_4Cl$, in the presence of light for pulse (3, 6, 12 and 36 h) and chase (3 and 12 h), and in the dark for 36 h. ag: albumen gland, dt: digestive tubule, fo: gonadal follicle.

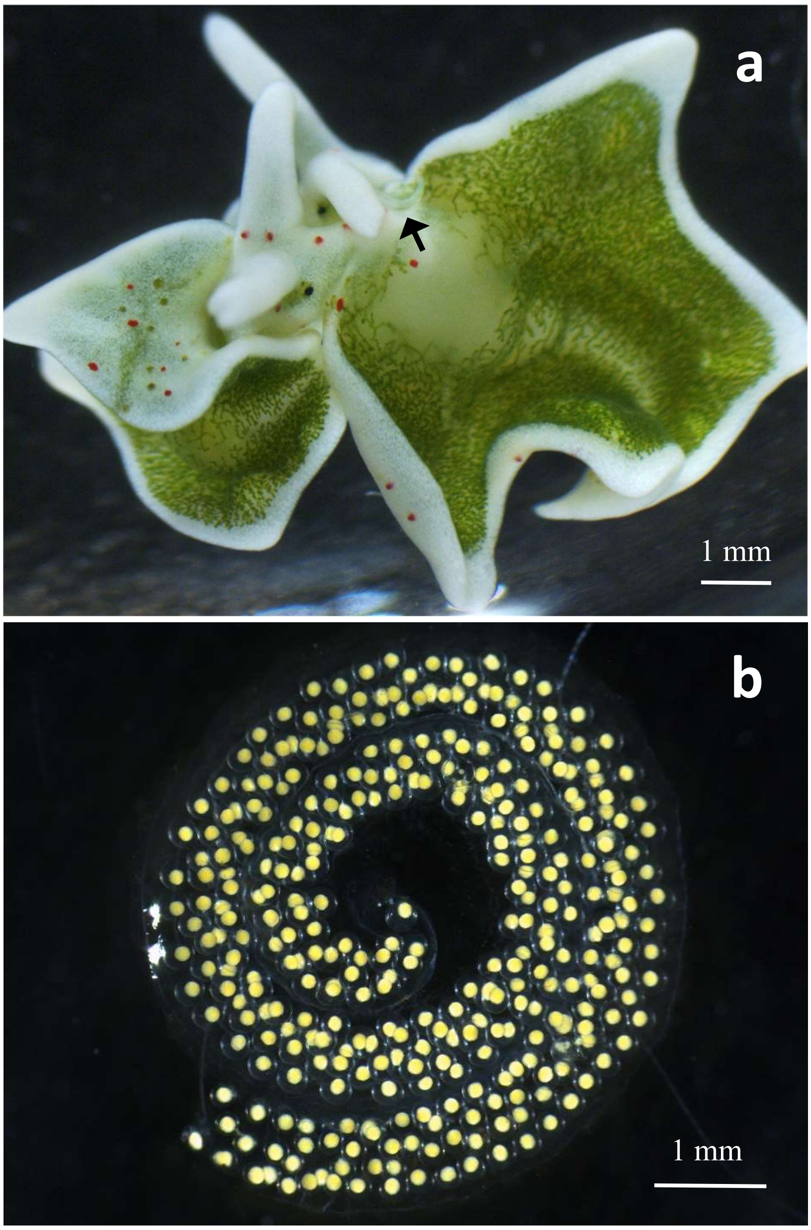


**Fig. S2.** ***Elysia timida*** **and spawned egg masses.** *E. timida* individuals taking up copulation position before reciprocal insemination (a). The black arrow signals the protruded penis. Spiral-shaped egg mass spawned by *E. timida* (b).

| | *Acetabularia acetabulum* | | *Elysia timida* | |
|---|---|---|---|---|
| **Fatty acid** | **Mean** | **SE** | **Mean** | **SE** |
| 14:0 | 0.71 | 0.05 | 0.32 | 0.03 |
| 16:0 | 19.68 | 1.06 | 14.82 | 0.61 |
| 18:0 | 10.39 | 1.93 | 10.80 | 0.94 |
| 20:0 | - | - | 0.17 | 0.02 |
| 24:0 | 0.10 | 0.03 | - | - |
| **Σ SFA** | **30.87** | **2.83** | **26.11** | **1.57** |
| 16:1n-9 | - | - | 0.13 | 0.04 |
| 16:1n-7 | 2.59 | 0.17 | 1.48 | 0.14 |
| 16:1 | 0.73 | 0.02 | 0.15 | 0.04 |
| 18:1 | - | - | 1.35 | 0.11 |
| 18:1n-9 | 7.89 | 0.39 | 8.67 | 0.35 |
| 18:1n-6 | 4.01 | 0.18 | 1.63 | 0.04 |
| 20:1n-11 | - | - | 2.34 | 0.14 |
| 20:1n-9 | 0.17 | 0.01 | 0.64 | 0.08 |
| **Σ MUFA** | **15.40** | **0.66** | **16.39** | **0.47** |
| 16:2 | 0.11 | 0.01 | - | - |
| 16:2n-4 | 3.86 | 0.29 | 0.77 | 0.03 |
| 18:2 | 0.23 | 0.01 | 0.24 | 0.02 |
| 18:2n-6 | 12.13 | 0.54 | 7.32 | 0.18 |
| $^{\Delta5,11}$ 20:2 | - | - | 0.86 | 0.09 |
| 20:2n-6 | - | - | 2.48 | 0.09 |
| 22:2 | - | - | 0.64 | 0.10 |
| 16:3 | 1.19 | 0.06 | - | - |
| 16:3n-4 | 5.01 | 0.33 | 1.20 | 0.06 |
| 18:3n-6 | 2.29 | 0.10 | 0.68 | 0.02 |
| 18:3n-4 | 0.12 | 0.01 | 0.18 | 0.02 |
| 18:3n-3 | 4.37 | 0.25 | 2.18 | 0.10 |
| 20:3n-6 | - | - | 0.23 | 0.05 |
| 20:3n-3 | - | - | 0.62 | 0.12 |
| 20:3n-4 | - | - | 0.55 | 0.13 |
| $^{\Delta7,13,16}$ 22:3 | - | - | 1.26 | 0.13 |
| 18:4n-4 | 0.22 | 0.01 | 0.32 | 0.02 |
| 18:4n-3 | 15.28 | 0.76 | 5.11 | 0.10 |
| 20:4n-6 | 0.26 | 0.02 | 6.15 | 0.26 |
| 20:4n-3 | 1.60 | 0.15 | 0.36 | 0.05 |
| 22:4n-6 | - | - | 10.13 | 0.44 |
| 20:5n-3 | 3.83 | 0.30 | 9.29 | 0.45 |
| 22:5n-3 | 3.21 | 0.38 | 6.94 | 0.35 |
| **Σ PUFA** | **53.72** | **2.18** | **57.49** | **1.18** |

**Table S1.** Fatty acid composition of *Elysia timida* and *Acetabularia acetabulum*. Relative abundance (%; mean ± SE, n = 5) of fatty acids in the sea slug *Elysia timida* and its macroalgal food source *Acetabularia acetabulum*. SFA – Saturated fatty acids; MUFA – Monounsaturated fatty acids; PUFA – Polyunsaturated fatty acids.

| | Pulse 6h | | Pulse 12h | | Pulse 24h | | Pulse 36h | | Chase 3h | | Chase 12h | | Control Dark 36h | | Control Light $^{12}C$ | |
|---|---|---|---|---|---|---|---|---|---|---|---|---|---|---|---|---|
| **Fatty acid** | Mean | SE | Mean | SE | Mean | SE | Mean | SE | Mean | SE | Mean | SE | Mean | SE | Mean | SE |
| **14:0** | 408.0 | 90.0 | 424.2 | 68.0 | 1041.5 | 322.9 | 1841.3 | 310.0 | 1710.9 | 337.6 | 1604.9 | 328.3 | -13.0 | 3.0 | -26.5 | 3.4 |
| **16:0** | 268.1 | 38.3 | 328.6 | 41.2 | 936.9 | 114.6 | 1889.5 | 380.7 | 1797.1 | 428.8 | 1756.8 | 431.8 | -19.6 | 0.8 | -24.5 | 0.3 |
| **18:0** | 102.8 | 52.4 | 233.8 | 92.8 | 587.4 | 139.4 | 1254.9 | 130.8 | 2051.1 | 608.3 | 1366.9 | 165.9 | -15.3 | 2.1 | -23.3 | 1.5 |
| **20:0** | 246.8 | 30.2 | 312.0 | 0.0 | 374.7 | 173.7 | 1006.6 | 60.1 | 762.3 | 38.3 | 1726.7 | 738.9 | -19.3 | n.d. | -29.1 | 4.1 |
| **16:1n-7** | 183.3 | 59.9 | 170.8 | 88.5 | 370.1 | 41.1 | 929.7 | 244.4 | 727.6 | 230.1 | 668.3 | 162.6 | -24.0 | 0.2 | -24.1 | 1.1 |
| **18:1n-9** | 242.1 | 87.7 | 307.7 | 79.4 | 872.9 | 218.2 | 1884.9 | 414.1 | 1578.4 | 523.0 | 2047.5 | 525.5 | -24.5 | 0.2 | -26.1 | 0.5 |
| **20:1n-11** | 46.0 | 11.9 | 75.8 | 23.2 | 328.6 | 85.0 | 674.9 | 140.0 | 1098.1 | 256.8 | 885.1 | 58.1 | -17.0 | 2.5 | -22.2 | 3.5 |
| **20:1n-7** | 69.0 | 26.3 | 60.7 | 19.9 | 180.2 | 20.4 | 647.6 | n.d. | 454.2 | 276.9 | 1430.3 | 457.5 | -20.9 | 3.2 | -22.2 | 6.2 |
| **16:2n-4** | -22.2 | 4.3 | -21.1 | 5.3 | 3.1 | 11.0 | 10.1 | 20.8 | 34.1 | 42.6 | 41.6 | 10.2 | -27.4 | 1.4 | -28.3 | 0.3 |
| **18:2n-9** | 69.1 | 14.7 | 178.9 | 80.5 | 381.4 | 78.5 | 647.4 | 218.9 | 890.1 | 110.4 | 1338.1 | n.d. | -23.3 | 2.6 | -28.4 | 0.9 |
| **18:2n-6** | -13.7 | 6.5 | 4.7 | 10.9 | 66.7 | 26.3 | 142.7 | 29.4 | 228.1 | 57.7 | 275.8 | 27.7 | -27.1 | 0.2 | -27.4 | 0.5 |
| **18:2n-3** | 22.9 | 3.5 | 52.5 | 14.0 | 68.8 | 30.4 | 210.7 | 45.9 | 331.6 | 145.5 | 79.3 | 9.5 | -18.6 | 1.9 | -22.5 | 0.4 |
| **20:2n-9** | 56.9 | 22.2 | 101.8 | 54.9 | 548.3 | 99.1 | 889.1 | 215.0 | 1661.6 | 770.3 | 1384.7 | 397.7 | -23.4 | 1.0 | -22.4 | 7.4 |
| **20:2n-6** | 22.2 | 7.2 | 37.1 | 10.9 | 81.1 | 24.3 | 191.4 | 14.3 | 224.3 | 56.3 | 256.8 | 36.2 | -18.7 | 1.3 | -25.4 | 0.4 |
| **22:2n-9** | 23.4 | 7.6 | 59.4 | 28.9 | 453.7 | 55.2 | 826.7 | 280.9 | 519.0 | 273.2 | 919.8 | 33.1 | -22.6 | 1.2 | -27.0 | 1.1 |
| **16:3n-3** | -12.7 | 7.5 | 2.7 | 9.2 | 66.5 | 37.3 | 65.3 | 25.7 | 64.5 | 25.9 | 110.3 | 11.4 | -26.5 | 0.7 | -27.1 | 1.9 |
| **18:3n-6** | -32.2 | 5.5 | -18.1 | 6.6 | -12.1 | 17.3 | -38.4 | 27.1 | -73.9 | 27.2 | -182.6 | n.d. | -24.2 | 0.2 | -27.4 | 0.8 |
| **18:3n-3** | -30.2 | 0.4 | -18.0 | 8.4 | -6.3 | 5.6 | -6.6 | 6.8 | 9.9 | 16.7 | 25.4 | 10.6 | -24.8 | 1.0 | -27.7 | 0.4 |
| **20:3n-9** | -5.6 | 1.2 | 3.3 | 12.0 | 22.8 | 14.1 | 86.7 | 12.5 | 126.6 | 55.3 | 118.6 | 42.5 | -23.2 | 0.3 | -27.0 | 0.5 |
| **20:3n-6** | -4.9 | 8.5 | -0.5 | 26.1 | 11.7 | 19.0 | 52.4 | 5.4 | 192.4 | 117.3 | 120.7 | 24.0 | -20.6 | 0.6 | -26.3 | 0.5 |
| **18:4n-3** | -31.6 | 1.6 | -27.1 | 2.1 | -29.6 | 1.9 | -33.2 | 1.5 | -51.2 | 17.8 | -29.6 | 6.7 | -25.8 | 0.8 | -19.9 | 1.8 |
| **20:4n-6** | -21.2 | 1.3 | -16.1 | 5.5 | -1.7 | 5.0 | 23.9 | 8.5 | 39.3 | 17.9 | 32.5 | 6.1 | -24.6 | 0.1 | -27.2 | 1.3 |
| **20:4n-3** | 16.8 | 11.5 | 39.9 | 19.5 | 55.1 | 18.1 | 131.4 | 80.2 | 243.2 | 141.6 | 268.7 | 42.3 | -15.8 | 2.2 | -11.2 | 14.3 |
| **20:5n-3** | -22.8 | 1.4 | -17.7 | 3.7 | -8.3 | 4.2 | 5.4 | 10.2 | 34.3 | 19.6 | 20.1 | 25.6 | -25.5 | 0.3 | -26.3 | 0.6 |
| **22:4n-6** | -0.1 | 8.4 | 11.9 | 12.4 | 72.6 | 26.1 | 171.5 | 39.4 | 217.2 | 30.3 | 188.4 | 17.7 | -21.6 | 1.3 | -25.9 | 0.0 |
| **22:5n-3** | -16.5 | 2.7 | -6.7 | 4.5 | 25.8 | 13.3 | 83.2 | 19.7 | 141.9 | 40.3 | 129.0 | 24.7 | -20.2 | 0.7 | -24.2 | 0.1 |

**Table S2. $^{13}C$ incorporation in the fatty acids of *Elysia timida*.** $^{13}C$ (‰; mean ± SE, n = 3) in fatty acids of *E. timida* incubated in artificial seawater (ASW) enriched with 2 mM $NaH^{13}CO_3$ in the presence of light for 6, 12, 24 and 36 h (pulse phase) and after transfer to fresh ASW enriched in $NaH^{12}CO_3$ (chase phase) for 3 and 12 h. Control samples were dark-incubated for 36 h in ASW enriched with 2 mM $NaH^{13}CO_3$ and light-incubated for 36 h in ASW enriched with 2 mM $NaH^{12}CO_3$.

| | Actinic | | Non-actinic | |
|---|---|---|---|---|
| Fatty acid | Mean | SE | Mean | SE |
| 14:0 | 0.91 | 0.09 | 0.65 | 0.08 |
| 16:0 | 12.02 | 0.92 | 10.37 | 1.04 |
| 18:0 | 12.40 | 1.16 | 11.16 | 1.48 |
| 20:0 | 0.67 | 0.13 | 0.44 | 0.03 |
| **Σ SFA** | **26.01** | **1.77** | **22.61** | **2.61** |
| 16:1n-7 | 1.93 | 0.19 | 1.53 | 0.10 |
| 18:1 | 2.89 | 0.24 | 3.54 | 0.31 |
| 18:1n-9 | 9.77 | 0.42 | 10.05 | 0.43 |
| 18:1n-6 | 2.81 | 0.08 | 2.77 | 0.03 |
| 20:1n-11 | 1.79 | 0.12 | 1.66 | 0.15 |
| **Σ MUFA** | **19.19** | **0.67** | **19.56** | **0.87** |
| 16:2n-4 | 0.58 | 0.04 | 0.53 | 0.02 |
| 18:2 | 0.60 | 0.06 | 0.64 | 0.06 |
| 18:2n-6 | 6.01 | 0.33 | 6.53 | 0.24 |
| 18:2n-3 | 0.92 | 0.07 | 0.87 | 0.05 |
| $^{\Delta 5,11}$ 20:2 | 1.29 | 0.11 | 1.30 | 0.13 |
| 20:2n-6 | 1.91 | 0.08 | 2.12 | 0.11 |
| 18:3n-6 | 0.60 | 0.04 | 0.56 | 0.01 |
| 18:3n-4 | 0.65 | 0.03 | 0.57 | 0.03 |
| 18:3n-3 | 2.23 | 0.09 | 2.55 | 0.07 |
| $^{\Delta 5,11,14}$ 20:3 | 1.10 | 0.07 | 1.33 | 0.10 |
| 20:3n-6 | 0.60 | 0.05 | 0.56 | 0.04 |
| 20:3n-3 | 0.61 | 0.05 | 0.48 | 0.03 |
| $^{\Delta 7, 13, 16}$ 22:3 | 1.11 | 0.10 | 1.27 | 0.09 |
| 18:4n-4 | 0.69 | 0.03 | 0.60 | 0.03 |
| 18:4n-3 | 3.26 | 0.13 | 3.33 | 0.20 |
| 20:4n-6 | 5.26 | 0.28 | 5.42 | 0.20 |
| 20:4n-3 | 0.90 | 0.07 | 0.99 | 0.06 |
| 20:4n-1 | 0.60 | 0.06 | 0.46 | 0.03 |
| 22:4n-6 | 8.45 | 0.75 | 8.83 | 0.51 |
| 20:5n-3 | 9.11 | 0.26 | 9.95 | 0.51 |
| 22:5n-3 | 8.32 | 0.54 | 8.94 | 0.56 |
| **Σ PUFA** | **54.80** | **1.45** | **57.83** | **1.89** |

**Table S3. Fatty acid composition of *Elysia timida* eggs.** Relative abundance (%; mean ± SE, n = 6 or 5) of fatty acids in eggs spawned by *E. timida* subjected to a 14:10 h light/dark photoperiod and a scalar irradiance of 40-160 µmol photons $m^{-2}$ $s^{-1}$ (Actinic) or 5 µmol photons $m^{-2}$ $s^{-1}$ (Non-actinic) for 28 days. Egg masses analyzed were the last spawned by the couples of each experimental unit. Animals were fed continuously with *Acetabularia acetabulum*. SFA – Saturated fatty acids; MUFA – Monounsaturated fatty acids; PUFA – Polyunsaturated fatty acids.

| | Pulse phase | | | Chase phase | | Experimental purpose |
|---|---|---|---|---|---|---|
| | Treatment | | Sampling | Treatment | Sampling | |
| 1 | Light | Labelled-ASW | 6, 12, 24, 36 h | Control-ASW | 3, 12 h | Measurements of light-dependent carbon and nitrogen assimilation |
| 2 | | Control-ASW | 36 h | n.a. | | Control for NanoSIMS measurements of natural isotopic ratios (parameters $C_{nat}$ and $N_{nat}$; see Material and Methods) |
| 3 | Dark | Labelled-ASW | 36 h | n.a. | | Control for light-independent carbon and nitrogen assimilation |

**Table S4. Dual isotopic labelling incubations of *Elysia timida*.** Description of treatments and sampling points of an isotopic dual labelling pulse-chase experiment of *E. timida* incubated in labelled artificial sea water with 2 mM $NaH^{13}CO_3$ and 20 µM $^{15}NH_4Cl$ (labelled-ASW) and non-labelled artificial sea water (control-ASW); n.a. – not applicable.
